# Mutational synergy with *CREBBP* loss in lymphomagenesis identified through forward insertional mutagenesis in a new DLBCL mouse model

**DOI:** 10.1101/2024.03.25.586554

**Authors:** Nathalie Sakakini, Roy Straver, Dhoyazan M. A Azazi, Sarah J. Horton, Ryan Asby, Simon E. Richardson, Pedro Madrigal, Elizabeth J Soilleux, Rachael Bashford-Rogers, Jeroen de Ridder, Brian J. P Huntly

**Affiliations:** Department of Haematology, University of Cambridge, Cambridge, UK; Cambridge Stem Cell Institute, University of Cambridge, Cambridge, UK; Centre for Molecular Medicine, University Medical Centre, Utrecht University, the Netherlands; Department of Pathology, University of Cambridge, UK; Department of Biochemistry, University of Oxford, UK; Hematology Service, Cambridge University Hospitals, Cambridge, UK

## Abstract

Loss-of-function mutations in the gene encoding the acetyltransferase CREBBP have been reported in numerous cancers but are particularly frequent in lymphoid malignancies. However, the functional significance of CREBBP loss in transformation and disease progression, most likely through cooperation with secondary genetic hits, has not yet been fully unravelled. Similarly, the contribution of the initial cell population sustaining CREBBP loss in the course of disease remains elusive. Here, we developed a new lymphoma mouse model integrating *Crebbp* loss at various stages of B cell development with a transposon-based insertional mutagenesis system. We demonstrated that *Crebbp* loss from the HSPC compartment resulted in an aggressive DLBCL-like disease, recapitulating well-characterised histological and molecular features of the human disease, as well as the recently described enhanced CD24 expression. More importantly, we identified candidate genes functionally equivalent to patient mutated genes. Those genes, mainly related to B cell development and cellular signalling, may represent novel therapeutic targets. Overall, this new model provides a powerful resource in which to conduct future mechanistic and therapeutic studies.

## Introduction

The acetyltransferase CREBBP (Cyclic AMP Response Element Binding (CREB) binding protein, also called CBP and KAT3A) is a pivotal transcriptional regulator. Its effects on transcription are mediated at multiple levels. By acetylating histone proteins (including H3K18, K27, K56), it favours chromatin accessibility at active promoters and enhancers and facilitates transcription factor binding [1]. It also acetylates and modulates the activity of non-histone proteins, such as the tumour suppressor p53, whose transcriptional activity is enhanced by acetylation [2,3]. Finally, via its scaffolding properties it recruits members of the basal transcriptional machinery (TBP, TFIIB, RNA polymerase II) and other proteins to gene regulatory complexes to facilitate transcription [4].

CREBBP has a broad interactome (with more than 300 interactants identified), thus it is implicated in numerous biological processes. It is essential for embryonic development, as *Crebbp*-null mouse embryos die between E9.5-10.5 from severe anaemia and neurological defects [5]. CREBBP is also critical for the formation and maintenance of haematopoietic stem cells (HSC) [6,7].

Germline *CREBBP* loss-of-function mutations have been reported in Rubinstein-Taybi syndrome [8]. This condition confers a predisposition to cancer, suggesting that CREBBP has a tumour suppressor role. This has been confirmed by numerous studies describing inactivating somatic mutations in a number of solid tumours and haematological malignancies [9,10]. However, *CREBBP* mutations are particularly prevalent in lymphoid malignancies. They occur in up to 40% of diffuse large B-cell lymphoma (DLBCL) [11] and up to 60% of follicular lymphoma (FL) [12], the two most common lymphoma subtypes. These mutations are mainly monoallelic and result in a partial loss of acetyltransferase activity [11]. Evidence from mouse models [3,13,14] demonstrates that, although *Crebbp* loss of function constitutes an early event in lymphomagenesis, it is not sufficient to drive full malignant transformation. This suggests the acquisition of secondary mutations are required, and we have shown that this is, at least in part, due to a defective DNA damage response related to the failure of p53 acetylation by Crebbp [3].

Despite global efforts to elucidate the molecular basis of DLBCL, the identity of the initial cell that acquires *CREBBP* mutations and the dynamics with cooperating mutations driving lymphomagenesis still remain to be fully address. To address these questions, we generated a new DLBCL mouse model that combines *Crebbp* loss with a transposon-based insertional mutagenesis (IM) system. Through the usage of haematopoietic stage-specific promoters controlling a recombinase Cre, this model was constructed in two versions (referred as IM-Mx1 or IM-Cd19) offering the flexibility to investigate *Crebbp* loss at different stages during lymphoid development. Both models synergize with dynamic insertions of the GrOnc transposon within the B-cell lineage, allowing evaluation of the effects of stochastic genomic insertions on disease burden. With this advanced approach we have shown that early *Crebbp* loss from the haematopoietic stem and progenitor (HSPC) compartment considerably increased the penetrance of B-cell lymphomas and considerably accelerated disease progression. Detailed analyses of murine tissues uncovered an aberrant B220^low^ B cell population expressing a germinal centre-like transcriptional program. In contrast, the loss of *Crebbp* at a later stage during B cell ontogeny resulted a longer latency period prior to B-cell lymphoma development. Sequencing analyses of the neoplastic B220^low^ cells isolated from our lymphoma models mapped GrOnc insertions to several key genes for B-cell development including *Pax5* and *Ebf1*, as well as in targetable signalling pathways. Comparisons with DLBCL patient data revealed significant functional overlaps between both datasets, validating the relevance of our new mouse models to investigate the molecular basis of this disease. Taken together this work provides a rational framework to design future studies.

## Materials and Methods

### Mice

C57Bl6 Mx1*Crebbp*-/- (*Crebbp*Fl/Fl; Mx1Cre/wt) and C57Bl6 Cd19Crebbp-/- (*Crebbp*Fl/Fl; Cd19Cre/wt) have previously been described [3]. Both conditional *Crebbp* knockout strains were independently bred with individual components of a forward mutagenesis system as illustrated in Supp Fig 1A, B. This system utilizes the GrOnc transposon [15] along with a Piggy Bac (PB) transposase under the transcriptional control of a modified Vk promoter (Vk*) containing perfect substrate sequences for somatic hypermutation, leading to the restoration of frame expression of the transposase within the germinal centre (GC) B-cell compartment [16]. Cre-mediated recombination was induced by intraperitoneal injections of 5 doses of poly(I) poly(C) (pIpC) (400 ug per dose; Sigma) administered every other day for 10 days to 6- to 12-week-old mice, or occurred naturally upon Cd19 expression in the B cell lineage (Supp Fig 1C). Deletion efficiency was determined by PCR of genomic DNA extracted from various tissues (Supp Fig 1D). Somatic hypermutation and expression of the transposase was triggered by Sheep Red Blood Cells (SRBC) stimulation. Three injections of 1x10^6 cells were a administered fortnightly (Supp Fig 1C). Peripheral blood samples were taken from the lateral saphenous vein and collected in EDTA-treated tubes. Peripheral blood cells were counted using a Vet abc automated counter (Scil Vet abc Plus+, Horiba Medical). Mice were closely monitored for signs of disease. Necropsy was performed to assess the presence of abnormalities. All mice were housed in a pathogen-free animal facility. Mice were maintained in a 12h-12h dark-light cycle, at a temperature of 21 degrees and 40% humidity. Experiments were conducted under UK Home Office regulations (under the Animals (Scientific Procedures) Act 1986, Amendment Regulations (2012)) and following ethical review by the University of Cambridge Animal Welfare and Ethical Review Body.

### Flow cytometry analyses

Single-cell suspensions were obtained from various tissues. Cells were filtered through a 70um nylon cell strainer (EASYstrainer, Greiner) and treated with red blood cell lysis buffer (Thermo Scientific) prior to staining. All staining were performed in DPBS (Invitrogen) supplemented with 2% FBS (Sigma) for 20 min at 4°C. A list of the antibodies used for the flow cytometry analyses is shown in Supplemental Table 1. Dead cells were excluded by gating on 7AAD negative cells (BD Pharmingen). Unstained, single stained and Fluorescence Minus One (FMO) controls were used to determine the background, fix the compensation in each channel and to determine the gates. Flow cytometry experiments were acquired on a Fortessa LSR (BD Bioscience). Data were analysed with FlowJo v10.8.2 software.

### Histopathology

Tissues were fixed in a 10% neutral buffered formalin solution (CellPath Ltd.). They were washed in PBS then transferred to 70% ethanol and embedded in paraffin blocks. Tissue sections (4um) were stained with Hematoxylin and Eosin (Thermo Fisher Scientific) according to the manufacturer’s protocol. Scoring of the slides was performed by an experienced histopathologist blinded to mouse genotypes.

### Transplantation assay

Total splenocytes (1 x 10^6) isolated from individual malignant mice were injected intravenously into lethally irradiated (5.5Gy + 5Gy) recipient mice that constitutively express EGFP. Helper bone marrow cells (0.5 x 10^6) from the recipient strain were mixed with the donor cells prior to the injection. EGFP^+^ cells were gated out to assess engraftment.

### BCR amplification and library preparation

Total RNA was isolated from FACS-sorted B220^low^/CD19^+^ splenocytes from sick mice (B-cell lymphoma only) using an RNeasy Micro kit (Qiagen). Control RNA was isolated from normal B220^high^/CD19^+^ splenocytes from control healthy mice that had also received pIpC and SRBC injections similarly to the experimental mice. Mice were age-matched, and gender balanced between each cohort whenever possible. RNA quantity and quality were checked by Qubit and Tape Station. Samples with a RIN score > 7 were used for library preparation. 200ng mRNA was used for BCR-sequencing library preparation. Briefly, mRNA was first retrotranscribed using a pool of primers targeting the IgH mouse locus. cDNA was then PCR amplified with 1ul of VH forward primer set pools (group 1 or group 2, 10uM stock per primer) and 1ul of a universal constant reverse primer in two independent reactions, using 25ul of KAPA High Fidelity master mix (Roche). The following PCR program was used: 5 min at 95 °C; five cycles of 5 s at 98 °C and 2 min at 72 °C; five cycles of 5 s at 98 °C, 10 s at 65 °C, and 2 min at 72 °C; and 30 cycles of 20 s at 98 °C, 30 s at 60 °C, and 2 min at 72 °C; with a final extension cycle of 7 min at 72 °C. Group 1 and group 2 PCR products from each sample were pooled and purified using SPRI beads and eluted in 50ul water. Libraries were generated with a KAPA Hyper Kit (Roche), incorporating adaptors and index primers provided by NEBNext® Multiplex Oligos for Illumina® (96 Unique Dual Index Primer Pairs). Final libraries were quantified and multiplexed for paired-end 300bp sequencing on an Illumina MiSeq platform.

### BCR-sequencing data processing and analysis

Forward and reverse reads generated from NGS sequencing were merged together if they contained an identical overlapping region >7 bp, or otherwise discarded, using FLASHv1.2.11 [17]. Joined reads were filtered for base quality (median Phred score >32) using QUASR v.7.01 (http://sourceforge.net/projects/quasr) [18] and converted to fasta format. Universal barcoded regions were identified in reads and oriented to read from V-primer to constant region primer. The barcoded region within each primer was identified and checked for conserved bases. Primers and constant regions were trimmed from each sequence, and sequences were retained only if there was >80% per base sequence similarity between all sequences obtained with the same barcode, otherwise they were discarded. The constant region allele with the highest sequence similarity was identified by 10-mer matching to the reference constant region genes from the IMGT database [19], and sequences were trimmed to give only the region of the sequence corresponding to the variable (VDJ) regions. Isotype usage information for each BCR was retained throughout the analysis hereafter. Sequences without complete reading frames and non-immunoglobulin sequences were removed and only reads with significant similarity to reference IgHV and J genes from the IMGT database using BLAST (v2.10.0) [20] were retained. Immunoglobulin gene usages and sequence annotation were performed in IMGT V-QUEST (v.3.4.9) [19], and class-switch recombination analysis was performed as in Bashford-Rogers et al. [21]. Repertoire statistics were performed in R using MANOVA for significance.

The network generation algorithm and network properties were calculated as in Bashford-Rogers et al. [21]: each vertex represents a unique sequence, where relative vertex size is proportional to the number of identical reads. Edges join vertices that differ by single-nucleotide nonindel differences and clusters are collections of related, connected vertices. A clone (cluster) refers to a group of clonally related B cells, each containing BCRs with identical CDR3 regions and IgHV gene usage or differing by single point mutations. Each cluster should originate from the same pre-B cell.

### RNA-sequencing library preparation

Total RNA was isolated from FACS-sorted B220^low^/CD19^+^ splenocytes from sick mice (B-cell lymphoma only) using a RNeasy Micro kit (Qiagen). Control RNA was isolated from normal B220^high^/CD19^+^ splenocytes from control healthy mice that also received pIpC and SRBC injections similarly to the experimental mice. Mice were age-matched, and gender balanced between each cohort whenever possible. RNA quantity and quality were checked by Qubit and Tape Station. Samples with a RIN score > 7 were used for library preparation. RNA-sequencing libraries were prepared using the NEBNext® Ultra™ II Directional RNA Library Prep kit for Illumina® (NEB) following supplier’s recommendations. Poly(A) mRNA was first separated using magnetic beads (NEB) and subsequently converted to cDNA. After adaptor ligation, libraries were amplified and barcoded by PCR using NEBNext® Multiplex Oligos for Illumina® (96 Unique Dual Index Primer Pairs). Libraries were quantified and multiplexed for paired-end 50bp sequencing on the Illumina Novaseq platform.

### RNA-sequencing data processing and analysis

Paired-end RNA-sequencing sequences were first quality-assessed before any further processing using FastQC (https://www.bioinformatics.babraham.ac.uk/projects/fastqc/). RNA-sequencing libraries that were found to pass the minimum quality thresholds across all quality control metrics were then subjected to adapter sequence trimming using TrimGalore package (https://github.com/FelixKrueger/TrimGalore). This was followed by alignment to the mouse genome (mm10) using STAR version=2.7.10a [22]. These flags (NH:i:1 and MAPQ=255) were used to extract uniquely-mapping high-confidence reads from alignment files, and used for transcriptome quantification with featureCounts version 2.0.1 [23]. Only genes with a raw read count of >= 10 in at least two samples were considered for all downstream analysis. Differential expression analysis was carried out on normalised gene-level transcriptomic reads using the Bioconductor package DESeq2 [24]. The R package ‘clusterProfiler’ (version 4.4.4) was used to perform KEGG Pathway enrichment, Gene Set Enrichment Analyses (GSEA) and Gene Ontology (GO) analyses on statistically-significant differentially-expressed genes [25]. A false-discovery rate (FDR) or a multiple-testing adjusted p-value of 0.05 was used across all tests to establish statistical significance. Differentially expressed genes were filtered using an FDR < 5% and Log2 Fold Change >|1|.

### Prediction of murine immune cell type from RNA-sequencing data

Bulk RNA-sequencing profiles of murine haematopoietic cell populations of various lineages and multiple differentiation trajectories were obtained from published datasets (accession codes: GSE79672, GSE83436, GSE109125 and GSE133743) [26,27,28,29]. Original sequencing runs were downloaded and processed as described above in the RNA-sequencing analysis section, to produce raw sequencing counts per gene for all libraries. These reads were subsequently normalised to fragments per kilobase of transcript per million mapped reads (FPKM). The FPKM-normalised read counts of each cell type were used as a reference dataset to predict the differentiation status of our murine tumour cells using the R package SingleR (version 1.10.0) [30]. The “SingleR” function was used to return the best-fitting annotation for each mouse library in our contingent, using the labelled reference dataset with similarly-normalised read counts of the same features (genes).

### Transposon-sequencing library preparation

Only mice showing evidence of B-cell lymphoma at the time of death were analysed. Genomic DNA was isolated from total mouse splenocytes using PureLink Genomic DNA kit (Invitrogen). 500ng of DNA was subsequently used for library preparation using NEBNext® Ultra™ II FS DNA Library Prep Kit for Illumina® following manufacturer’s recommendations. Briefly, DNA was fragmented for 15 minutes at 37 C. After end repair and A tailing steps, splinkerette adaptors were ligated before PCR amplification using 2x KAPA HiFi HS ReadyMix kit. After purification with SPRI beads, libraries were amplified and barcoded by PCR using TraDIS indexes. DNA profiles were visually inspected using a Tape Station D5000. Final libraries were quantified by qPCR using Library Quant Standards (Illumina) and pooled in equimolar amount for paired-end 75bp sequencing on an Illumina MiSeq platform.

### Transposon-sequencing analysis and Common Insertion Sites identification

After sequencing, each sample existed across a set of 4 fastq files: One pair for “left handed” and one pair for “right handed” data. The pairs refer to the forward and reverse sequenced reads for each “handed” set. Files within a pair were processed together. Read pairs were filtered out from the fastq files where the first read did not start with the expected junction sequence or did contain the sequence of the non-mobilized transposon, Junction sequence from the first read in each pair was trimmed for all reads kept. The processed reads were then mapped (forward-reverse pairs combined) using *bwa mem*, followed by sorting and indexing using *samtools*. Read pairs where the first read was not mapped were rejected. The start location of each first read, indicating the exact site of an insertion, was retrieved. For further processing both “handedness” start sites were combined: all reads starting at the exact same base with the same mapping directionality compared to their “handedness” were treated together, creating (up to) two stacks of reads per insertion site: combining Left-Forward and Right-Reverse reads in one stack, and Left-Reverse and Right-Forward reads in the other stack, making each read stack specific to the direction of an insertion at that location. Reads in each stack were filtered based on mapping quality (>=20) and alignment score (>=30). Reads that did not meet these thresholds, and reads where the second in pair was mapped on another reference sequence or further than 1500 bp away, were removed. Each stack was given a non-duplicate score (*ndup*), indicating how many reads after filtering had the same mapped start position in the first read but a different mapped end position in the second read. This *ndup* count reduces the odds of double counting read duplicates and provides a diminishing returns effect for sites with extremely high numbers of reads. For each sample, the sum of all insertion sites’ *ndup* scores on all chromosomes other than chromosome 19 (origin of the transposon) was calculated and a cutoff was set at 17.5% of this summed value. All chromosomal mapped insertion sites (including chromosome 19) in the sample were then filtered, removing sites with an *ndup* value below the cutoff. This approach removes putative insertion sites for which there is relatively little evidence while adjusting for sequencing depth variations among samples. Next, the insertion sites of all samples were merged into a single file for further processing with Kernel Convolution Rule Based Method (KCRBM) [31,32]. We altered KCRBM to work with *BSgenome*.*Mmusculus*.*UCSC*.*mm10*, use the *PB* insertion system, enabled *Local Hopping Correction* (LHC) and used the *ndup* values as insertion depth per insertion site. The insertion sites output was then further processed and visualised using custom python scripts.

## Results

### Early Crebbp loss cooperates with insertional mutagenesis to significantly accelerate the B-cell lymphoma phenotype in a novel murine model

To delineate the relative contribution of the haematopoietic compartment which initially acquires Crebbp loss-of-function mutation and the effect of mutational synergy with *Crebbp*-loss in lymphoma development, we generated a new DLBCL mouse model by introducing a forward transpositional mutagenesis system into our existing *Crebbp* conditional knockout mouse model [3]. Briefly, Mx1*Crebbp*^Fl/Fl^ mice were independently crossed with either Vk* PB or GrOnc mice to generate the experimental Vk* PB^Tg/wt^; GrOnc^Tg/wt^; Mx1*Crebbp*^Fl/Fl^ mice (here after called IM-Mx1*Crebbp*) (Supp Fig 1A, B). Cre-mediated *Crebbp* excision initially occurs in the HSPC compartment and is propagated in subsequent compartments, where it combines with transposition of the GrOnc transposon in a B-cell lineage restricted manner. In keeping with the requirement for further mutations to manifest full lymphoma transformation, this new model considerably increased the penetrance of B-cell disease, where it accounted for ∼80% of all deaths (Fig1A). In comparison with IM-Mx1*Crebbp*^+/+^ littermates that also had B-cell restricted insertional mutagenesis, mice with early loss of *Crebbp* displayed a significantly shorter latency (median survival 12.3 vs 17.8 weeks, p<0.05) and a slightly higher proportion of B-cell diseases (Fig 1A, B). Macroscopically, mice from both genotypes presented with hepatosplenomegaly and elevated white blood cell count (WCC) (Fig 1C, Supp Fig 1E, F) although histological features were distinct for *Crebbp* deleted and WT cases. Lymphadenopathy was also observed in both models, however this was predominantly a feature of the IM-Mx1*Crebbp*^-/-^ mice (referred as ‘nodal’) (Fig 1D). Extranodal disease infiltration was also seen in both genotypes but was more common in the IM-Mx1*Crebpb*^*+/+*^ mice (referred as ‘extranodal’, Fig 1D). Histological examination revealed disruption of normal tissue architecture of the spleen and the lymph nodes and extension infiltration by neoplastic cells. Extensive infiltration spread to other tissues including liver and kidneys (Fig 1D, E), consistent with an aggressive DLBCL-like disease and previously described DLBCL mouse models [33]. The lymphomas were most commonly classified as DLBCL for both genotypes, with a higher proportion in the IM-Mx1Crebbp^-/-^ cohort (64.5% vs 53.8% (Fig 1F) rising to 87.1% vs 69.2% when considering cases showing a mixture of DLBCL and FL). Conversely, the incidence of BL was increased in the IM-Mx1-*Crebbp*^+/+^ cohort (19.2% vs 3.2%) (Fig 1F). These results are consistent with observations in patients, where *CREBBP* mutations are more frequently observed in DLBCL and FL, and are much less common in BL. Finally, the disease efficiently transplanted to recipient mice (Fig 1G) and recapitulated features of the initial disease, including hepatosplenomegaly (Supp Fig 1G, H).

**Figure 1:**
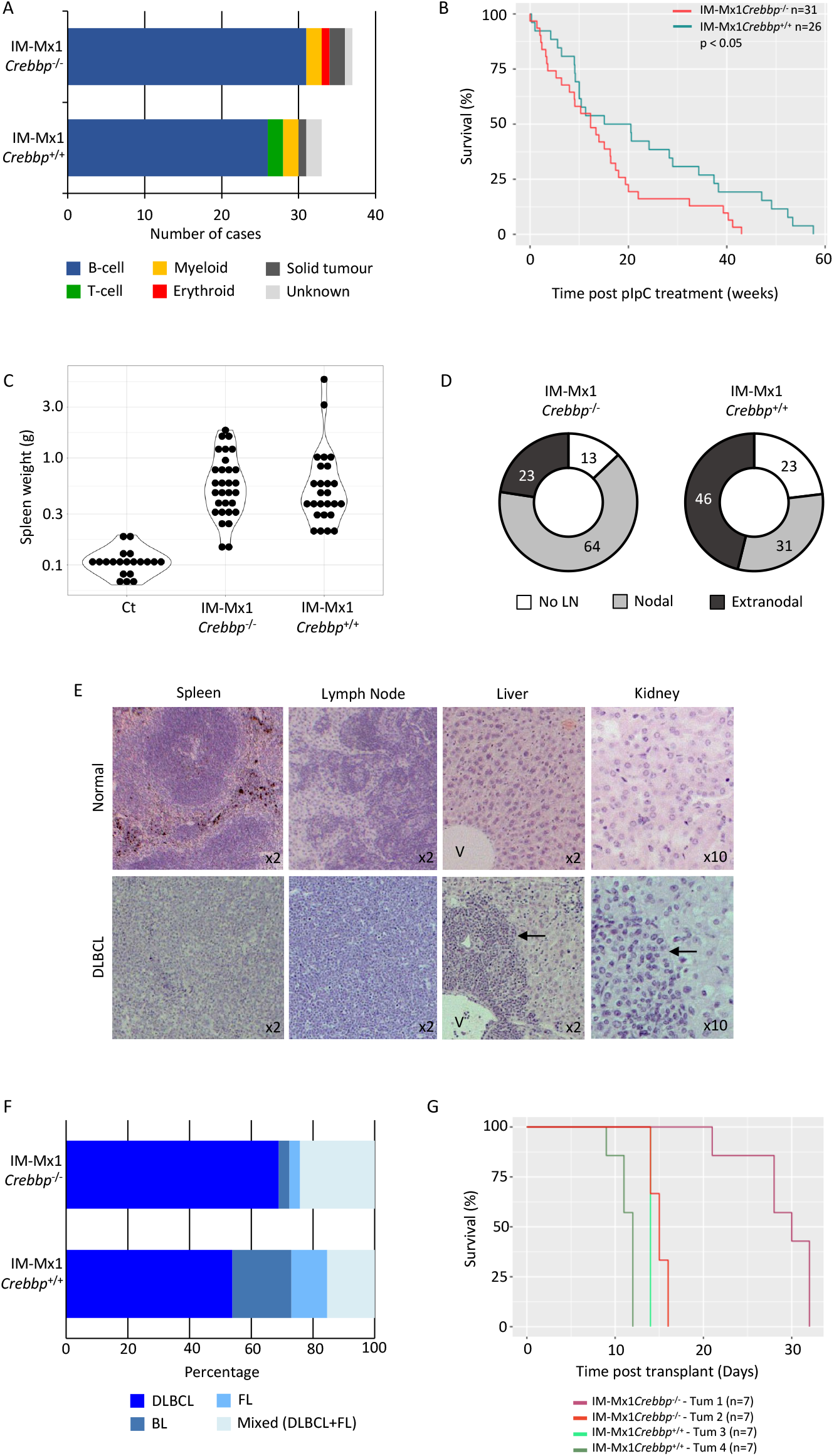
Early Crebbp loss cooperates with insertional mutagenesis to significantly accelerate the B-cell lymphoma phenotype in a novel murine model. (A). Bar chart showing a high incidence of B-cell malignancies in both models. (B). Survival curve demonstrating a significantly shorter survival in IM-Mx1*Crebbp*^-/-^ mice (n=31) comparted to IM-Mx1*Crebbp*^+/+^ littermates (n=26). Only mice with evidence of B-cell malignancies were considered in this analysis. P<0.05, Log Rank test. (C). Violin plot showing the spleen weight of lymphoma mice compared to control wild-type mice. Each dot represents a mouse. (D). Pie chart comparing tissue infiltration between both lymphoma models. Mice showing enlarged spleen and liver only were scored as “No lymph node” (white), when they also had enlarged lymph nodes as “Nodal” (grey), when they had evidence of additional tissue infiltration with or without lymph node involvement were scored as “Extranodal” (dark grey). (E). Hematoxylin and eosin staining of mouse tissue sections comparing healthy *versus* DLBCL-like disease. (F). Bar chart showing the distribution of B cell lymphoma subtypes in both mouse models based on histological features. Mixed cases refer to mice with evidence of DLBCL and FL. (G). Survival curve demonstrating an extremely short survival after primary tumour transplant into recipient mice. Four primary tumours of indicated genotypes (coloured lines) were transplanted into 7 recipient mice each. LN, lymph node, V, hepatic vein, DLBCL, diffuse large B cell lymphoma, BL, Burkitt lymphoma, FL, Follicular lymphoma.

### An aberrant B220^low^/CD19^+^ population drives a DLBCL-like disease

To further characterise our new model, we analysed single cell suspensions from lymphoma mouse spleens to determine their immunophenotype. These analyses revealed the presence of an expanded and aberrant B220^low^/CD19^+^ population which was completely absent from the spleen of control animals (Fig 2A). These neoplastic cells were also demonstrated across all tissues analysed, consistent with the histological tissue infiltration (Fig 2B). The cells expressed PNA, GL7 and, to a lesser extend, Fas, reminiscent of a germinal centre phenotype (Fig 2C, Supp Fig 2A). Interestingly, they also demonstrated enhanced CD24 expression (Fig 2D, Supp Fig 2B). These cells were capable of reconstituting and giving rise to disease in recipient mice after transplant, and the tumours that developed retained the same immunophenotype as the primary DLBCL-like cells (Supp Fig 2C). Deep sequencing analysis of B cell receptor repertoires was performed across lymphomas from IM-Mx1*Crebbp*^-/-^, IM-Mx1*Crebbp*^+/+^, assessing B220^low^/CD19^+^ cells as tumour cells and compared to B220^high^/CD19^+^ cells from control mice. This revealed three main features. All tumours demonstrated BCR VDJ rearrangements, with evidence of low levels of class switch recombination and modest somatic hypermutation in keeping with their mature B cell origin (Fig 2E, F). Multiple expanded clones per mouse were observed, however there was typically a single large expansion within the IM-Mx1*Crebbp*^-/-^ tumours, suggesting that the absence of Crebbp alters clonal dynamics during lymphomagenesis (Fig 2G, H). Moreover, the consensus sequences of the largest 3 clones per mouse were almost identical to the germline, supportive of antigen-inexperienced B cells as precursors of these lymphomas (Supp Table 2). Expanded clones were predominantly IgHM with no differences in isotype usages between clones of different genotypes, however, the relative proportion of class-switch recombination from IGHD/M to IGHG2B and IGHA was significantly higher in IM-Mx1*Crebbp*^-/-^ tumours than IM-Mx1*Crebbp*^+/+^ and controls (Fig 2H).

**Figure 2:**
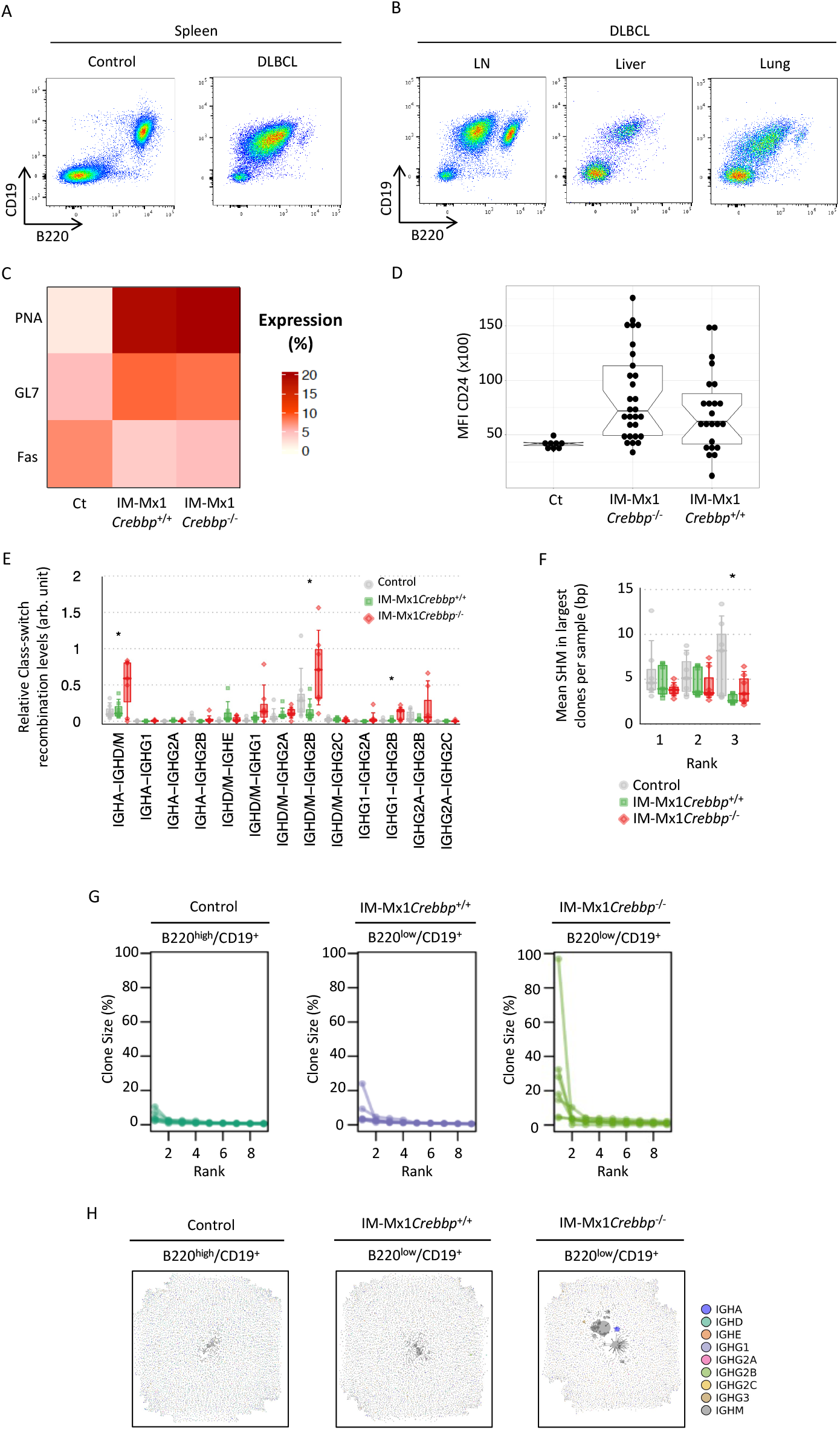
An aberrant B220^low^/CD19^+^ population drives a DLBCL-like disease. (A). FACS plot showing B220 and CD19 cell surface marker expression in the spleen comparing DLBCL-like lymphoma versus control. (B). FACS plot showing B220 and CD19 cell surface marker expression in DLBCL-like lymphoma mice across multiple tissues. (C). Heat map showing the median expression of germinal centre markers in the neoplastic B220^low^/CD19^+^ population from both lymphoma models in comparison to control B220^high^/CD19^+^ splenocytes. (D). Boxplot showing the Mean Fluorescence Intensity (MFI) of CD24 in the neoplastic B220^low^/CD19^+^ population from both lymphoma models in comparison to control B220^high^/CD19^+^ splenocytes. (E). Graph showing the relative level of class-switch recombination per indicated population. (F). Graph showing the mean number of somatic hypermutations (SMH) in the three largest clones per indicated population. (G). Graphs showing the clone size (as percentage) of the top largest clones per indicated population isolated from control or lymphoma mice. (H) Representative BCR network plots of deep sequenced PCR amplified immunoglobulin variable gene regions of indicated populations isolated from control mice (n=7) or primary tumour samples (n=5-7).

### Transcriptomic alterations in lymphoma tumours mimic changes in DLBCL patients

To define similarities between our new mouse models and molecular features of human DLBCL, we performed bulk RNA-sequencing on sorted B220^low^/CD19^+^ splenocytes from both lymphoma models and compared them to control B220^high^/CD19^+^ splenocytes from wild-type animals. Comparisons with the control revealed profound transcriptional changes in the lymphoma cells with nearly 7000 differentially expressed genes (6885 in IM-Mx1*Crebbp*^-/-^ vs Ct and 6620 in IM-Mx1*Crebbp*^+/+^ vs Ct) (Fig 3A). Approximatively 60% were commonly dysregulated in both models (Fig 3A). Amongst them the lymphoma-associated oncogenes *Ezh2* and *Mycn* [34] were up-regulated, alongside general proliferation genes including *E2f1, Cdk1, Cccnd3* and *Mki67* reflective of an increased cell cycle status. Down-regulated genes included several mediators of the immune response, such as *Cd40, Cd74* and the TNF family members *Tnfsf14* and *Tnfsf18*. In addition, class II MHC genes *Ciita, H2-Ab1, H2-Ob* and *H2-Eb2* previously demonstrated to be down-regulated in *Crebbp* mutated lymphoma [14,35] (Fig 3B) were also reduced in their expression. GSEA analysis comparing both lymphoma models found Crebbp-related signatures enriched in *Crebbp* replete mice, validating loss of Crebbp transcriptional activity after Cre-mediated excision in our IM-Mx1*Crebbp*^-/-^ mice (Fig 3C). Functional annotations showed that up-regulated genes in lymphoma vs control were significantly enriched for biological processes related to cell proliferation and cell cycle, whereas down-regulated genes were involved in immune response and other immune-related processes (Fig 3D, E). Quantitative and qualitative differences were also observed between lymphoma genotypes (Fig 3D). Using transcriptomic profiles available for normal mouse immune cell types [ImmGen Consortium,26,27,28,29], we were able to demonstrate that the B220^low^/CD19^+^ neoplastic population from both models closely resembles germinal centre cells at the transcriptomic level (Fig 3F).

**Figure 3:**
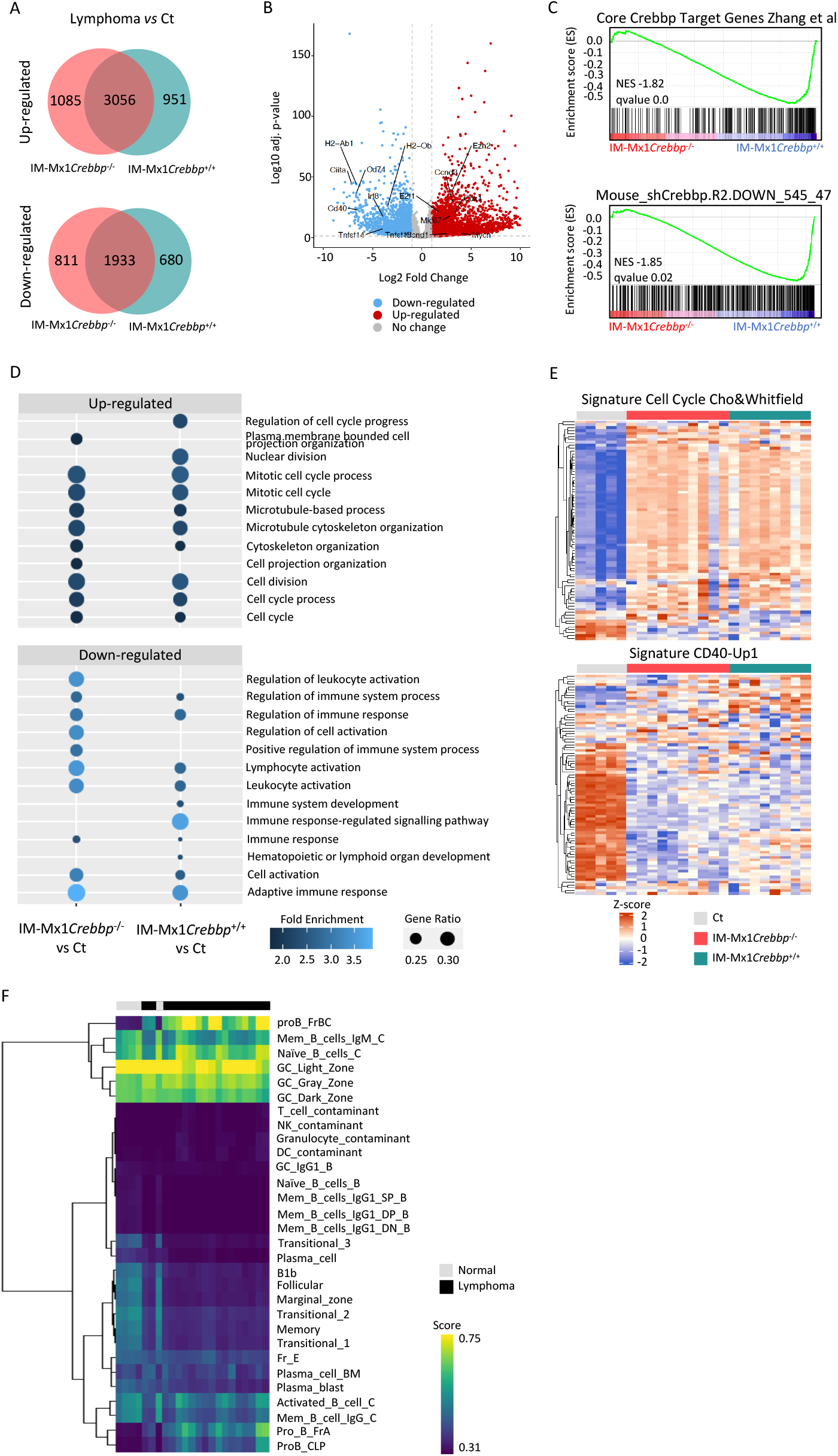
Transcriptomic alterations in lymphoma tumours mimic changes in DLBCL patients. (A). Venn diagram showing the overlap between the differentially expressed genes identified from lymphoma versus control comparisons. The numbers indicate the number of differentially expressed genes. (B). Volcano plot showing genes significantly dysregulated (FDR 5% & Log2FC >|1|) in IM-Mx1*Crebbp*^-/-^ vs control. (C). GSEA analysis showing two independent Crebbp signatures enriched in IM-Mx1*Crebbp*^+/+^. (D). Dot plot showing biological processes associated with up-regulated (top) and down-regulated (bottom) genes. (E). Selected heat maps illustrating the up-regulated and down-regulated processes. (F). Heat map showing the score of each sample analysed by RNA-sequencing for individual immune cell type provided by the Immunological Genome Project (ImmGen) and additional studies., BM, bone marrow, CLP, Common lymphoid progenitor, DC, Dendritic cell, GC, germinal centre, Mem, Memory.

### Loss of CREBBP later in B-cell development attenuates DLBCL-like disease

Our results show that *Crebbp* loss in early hematopoietic progenitors in association with forward mutagenesis drives an aggressive DLBCL-like phenotype. We have previously shown that losing Crebbp function later during lymphoid development significantly decreased lymphoma development [3] and thus we sought to test whether combination with insertional mutagenesis would overcome this delayed latency. To test this hypothesis, we generated a second mouse model that similarly allowed for B-cell transposition, where *Crebbp* excision occurred in a committed B-cell progenitor (hereafter called IM-Cd19*Crebbp*^-/-^). In this new combination, the majority of mice also develop B-cell lymphomas (75% vs 83.7% for IM-Mx1*Crebbp*^-/-^), of which about half the cases displayed unequivocal features of DLBCL, whereas the other half also demonstrated histological elements of FL (Fig 4A, B). The lower penetrance of DLBCL and the increased proportion of FL suggested that Crebbp loss-of-function within B-cell progenitors plus insertional mutagenesis leads to a less aggressive disease, perhaps more in keeping with transformed FL. This was consistent with the longer latency in IM-Cd19*Crebbp*^-/-^ mice (median survival 12.3 IM-Mx1*Crebbp*^-/-^ vs 51.9 weeks IM-Cd19*Crebbp*^-/-^) (Fig 4C). However, at their terminal end point the IM-Cd19*Crebbp*^-/-^ mice were almost undistinguishable from their Mx1 counterparts. Apart from a reduced nodal involvement, they presented with similarly enlarged spleens and livers, and elevated WCC (Fig 4D, E, Supp Fig 3A, B). At the cellular level, the neoplastic B220^low^/CD19^+^ population identified in our initial model was also detected in multiple tissues including the spleen, the lymph nodes and the liver (Fig 4F, Supp Fig 3C) and expressed the germinal centre markers PNA, GL7 and Fas (Fig 4G, Supp Fig 3D). Finally, we could also demonstrate that IM-Cd19*Crebbp*^-/-^ tumours were transplantable and generated disease in recipient mice (Fig 4H, Supp Fig 3E-G).

**Figure 4:**
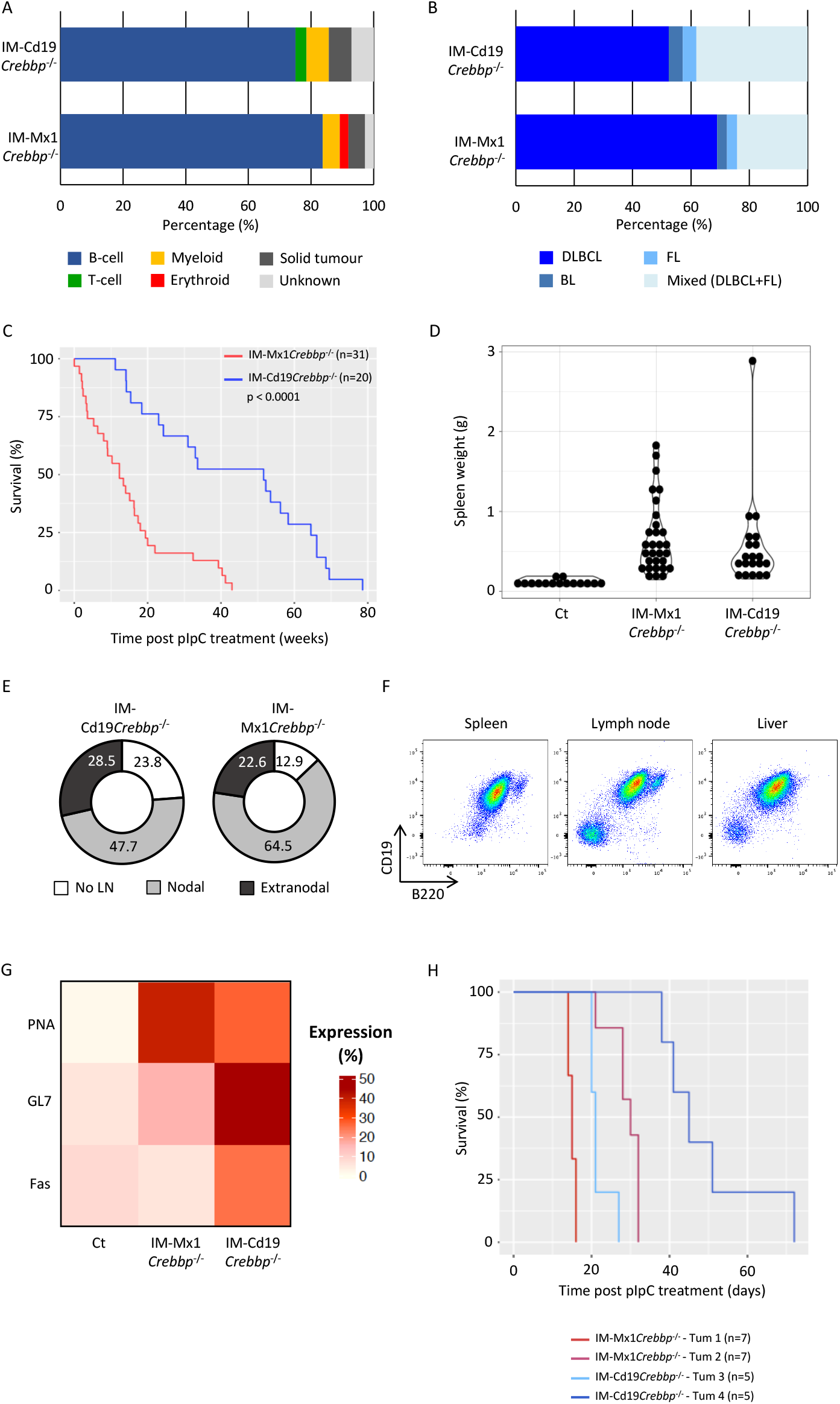
Loss of CREBBP in B-cell progenitors attenuates DLBCL-like disease. (A). Bar chart showing a high incidence of B-cell malignancies in both the IM-Cd19*Crebbp*^-/-^ and IM-Mx1 *Crebbp*^-/-^ models. (B). Bar chart showing the distribution of B-cell lymphoma subtypes in both mouse models based on histological features. Mixed cases refer to mice with evidence of DLBCL and FL. (C). Survival curve demonstrating a significantly longer survival in IM-Cd19*Crebbp*^-/-^ mice (n=20) comparted with IM-Mx1*Crebbp*^-/-^ mice (n=31). Only mice with evidence of B-cell malignancies were considered in this analysis. P<0.001, Log Rank test. (D). Violin plot showing the spleen weight of mice with lymphoma compared to control wild-type mice. Each dot represents a mouse. (E). Pie chart comparing tissue infiltration between both lymphoma models. Mice showing enlarged spleen and liver only were scored as “No lymph node” (white), when they also had enlarged lymph nodes as “Nodal” (grey), when they had evidence of additional tissue infiltration with or without lymph node involvement as “Extranodal” (dark grey). (F). FACS plot showing B220 and CD19 cell surface marker expression across multiple tissues isolated from IM-Cd19*Crebbpp*^-/-^mice. (G). Heat map showing the average expression of germinal centre markers in the neoplastic B220^low^/CD19^+^ population from IM-Mx1*Crebbp*^-/-^ and IM-Cd19*Crebbp*^-/-^ lymphoma mice in comparison to control B220^high^/CD19^+^ splenocytes from control mice. (H). Survival curve of recipient mice transplanted with primary tumours of indicated genotype (n=5-7 recipients). DLBCL, diffuse large B cell lymphoma, BL, Burkitt lymphoma, FL, Follicular lymphoma.

### Transposon Common insertional Sites (CIS) identify known and putative lymphoma oncogenes and tumour suppressor genes

To assess the contribution of cooperating mutations to the development of DLBCL, our mouse models utilised a transposition system that allows the cooperating lesions to be identified through isolation of recurrent insertions. We then sequenced and analysed the insertion sites in the lymphomas from 71 out of the 77 mice that presented with evidence of B cell lymphoma at the time of death. The 6 remaining lymphoma samples failed to pass quality control steps and were excluded from subsequent analysis. Ligation-mediated PCR followed by next generation sequencing from individual tumours generated over 34 million reads [36,37]. Of these reads, 91% were successfully paired and mapped to the mouse genome (version mm39). Using a Kernel Convolution Rule Based Method (KCRBM), we identified 1615 unique genomic insertions across all tumours [31,32], of which the majority were genotype specific (Fig 5A). When linked to the nearest gene, these insertions collapsed to 1264 genes, of which 87 genes harboured at least 3 independent insertional events (Fig 5B, Suppl Table 3). Amongst the most common targeted genes we found major regulators of B-cell development (*Ebf1, Pax5, Ikzf1*), alongside with well-known proto-oncogenes, particularly related to tyrosine kinase pathways (*Flt3, Stat5b, Nras* and *Jak2*) and known tumour suppressors (*Cdkn2a, Pten*) (Fig 5B, Supp Table 3). Most of these were organised in a highly connected network, suggesting their involvement in combinatorially coordinating the same processes (Fig 5C). Functional annotations supported this finding, showing significant enrichment of biological processes related to haematopoietic differentiation (interestingly both lymphoid and myeloid) and signal transduction through tyrosine kinases (Fig 5D). Additionally, many of the genes, such as *Nras, Flt3* and *Sos1*, are known for their contribution to several cancers, including haematological malignancies (Supp Fig 4A).

**Figure 5:**
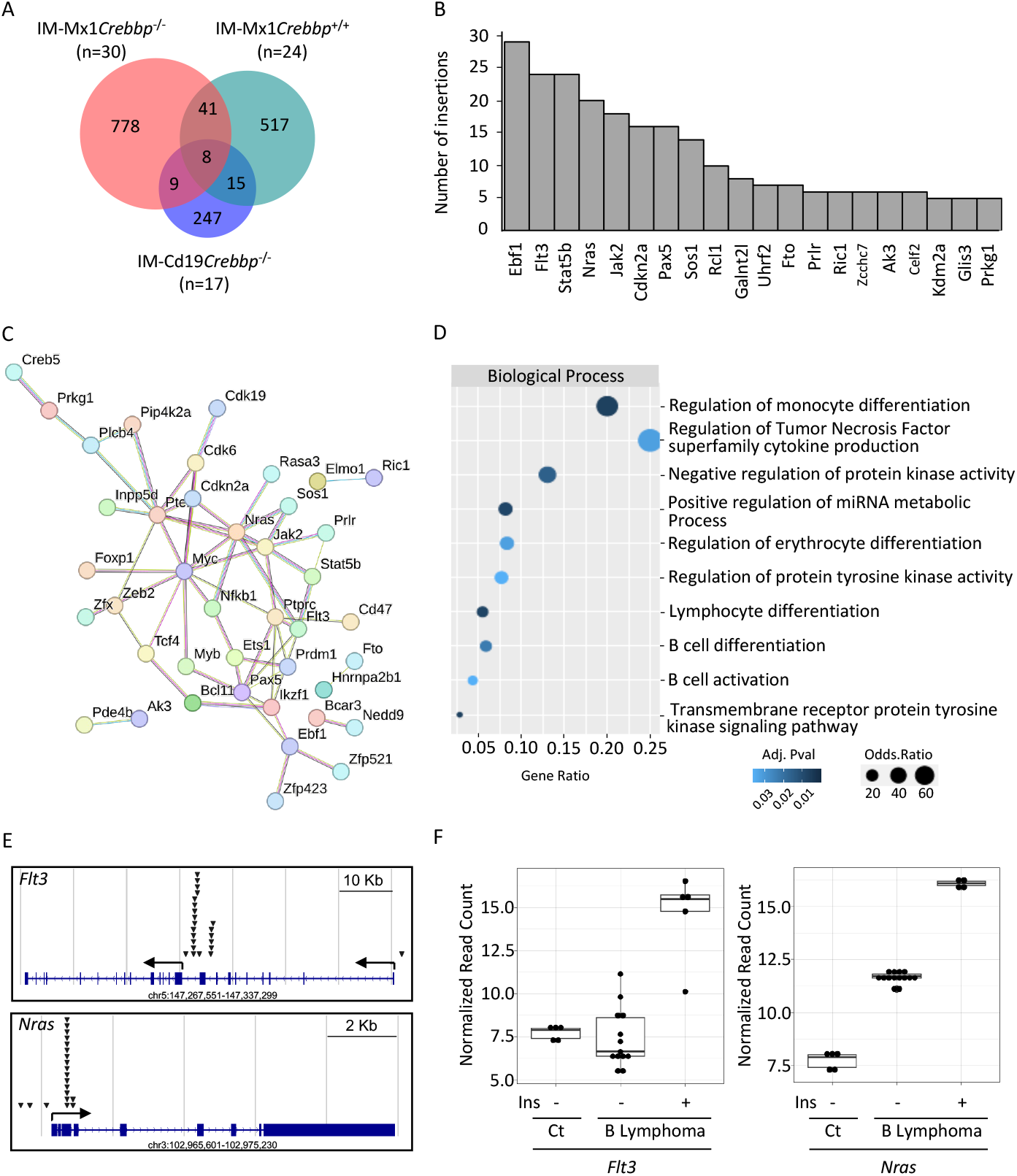
Identification of transposon common integration sites (CIS) (A). Venn diagram showing the distribution of individual insertions across genotypes. (B). Barplot showing the number of insertions for the top 20 most targeted genes. (C). String.db network showing interactions between the genes harbouring at least three insertions. Single nodes are not displayed. Interaction score confidence 0.6. (D). Functional annotations showing the Biological Processes for the genes insertionally targeted in mouse lymphoma models. (E). Screen shots showing the genomic locations of insertions for *Flt3* (top) and *Nras* (bottom) genomic loci. (F). Boxplots showing the normalised read count for *Flt3* (left) and *Nras* (right) in control (Ct) and B lymphoma tissues in presence (Ins +) or absence (Ins -) of the transposon GrOnc. Each dot represents an individual sample.

Because of its genetic design, the GrOnc transposon can alter gene expression through multiple mechanisms. It contains both strong promoter and splice donor, acceptor and bi-directional polyadenylation signals sequences. These can enhance gene expression when landing upstream of a transcriptional start site or conversely insertions occurring in introns can generate truncated proteins, leading to loss-of-function mutations (Supp Fig 1A) [38,37]. Knowing the genomic location of the insertions, it is therefore possible to predict their effect on neighbouring genes. We mapped the location of all the insertions and experimentally validated their effect on selected candidates. These included *Flt3* and *Nras*, where insertions occurred in a hotspot either upstream of a predicted transcriptional start site, or in the 5’ portion of the gene, leading to upregulation compared to tumours negative for those particular insertions (Fig 5E, F).

### Positive correlation between murine models and human data

Our new lymphoma mouse models faithfully recapitulated critical features of human DLBCL, such as nodal involvement. They also identified cooperating mutations involving genes important for normal B cell development such as *Ebf1* and *Pax5*, as well as known oncogenes and tumour suppressors such as *Ezh2* and *Cdkn2a*, respectively, which may be relevant for DLBCL progression. Analysis of gene-set libraries curated in the EnrichR tool [39,40] revealed that our candidate list of DLBCL cooperating mutations highly correlated with proteomic signatures of human lymphoid cell lines contained in CCLE proteomic database [41]. Out of the 17 significantly enriched proteomic profiles, 7 (41%) were from lymphoid cell lines, amongst which 3 were DLBCL lines (SUDHL4, OCI-Ly3, NUDHL1) (Fig 6A). We then investigated whether the genes mutated in our DLBCL mouse models were also mutated in DLBCL patients [42,43, 44,45]. 19/77 (∼25%) of our candidate genes that had a human ortholog were found to be mutated in DLBCL patients (Fig 6B). To get a more comprehensive view of the commonalities between both data sets, we performed a comparative functional analysis using Metascape [46]. We found a significant overlap between GO terms that were enriched in each data set, as exemplified by the B cell receptor signalling pathway and the process of leukocyte differentiation (Fig 6C, Supp Fig 5B). We also found a striking enrichment for gene-disease signatures from the DisGeNET database [47] related to various B cell lymphomas, amongst them diffuse large B-cell lymphoma and other lymphoid malignancies (Fig 6D). Given the strong interconnection between both datasets, we integrated them in one single protein interaction network, using Cytoscape. Interestingly this analysis highlighted several interactions between the genes mutated in our mouse model and those mutated in DLBCL patients such as JAK2 and NRAS (Fig 6E), suggesting that alterations of these signaling pathways are important in DLBCL pathogenesis.

**Figure 6:**
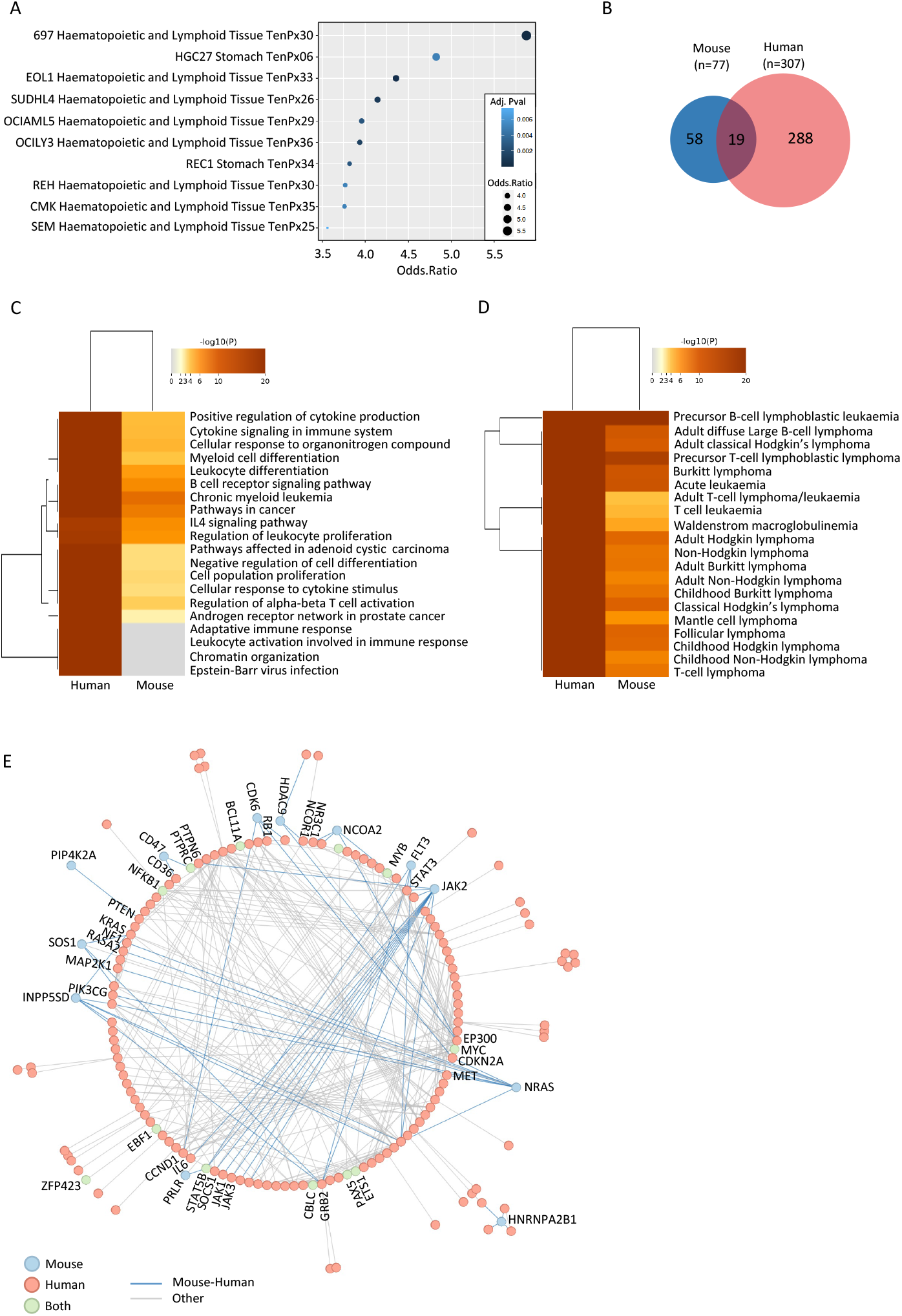
Positive correlation of murine models with human data. (A). Functional annotations showing significant enrichment of insertionally targeted genes in proteomic profiles of lymphoid cell lines contained in the CCLE proteomic database, using the EnrichR tool. (B). Venn diagram showing the overlap between mouse candidates (human orthologs) and the most recurrently mutated genes in DLBCL patients. (C). Heatmap showing comparative enrichments of GO terms for the genes identified from our insertional mouse screens and the most recurrently mutated genes in DLBCL patients. (D). Heatmap showing comparative enrichments of gene-disease signatures from the DisGeNET database, as designated in C. (E). Protein network showing interactions between genes as designated in C. Nodes are colour-coded according to their gene set of origin (blue for mouse, red for human, green when found in both). Edges represent interactions between nodes (blue for interactions between a mouse and a human gene, grey for any other combination).

## Discussion

Non Hodgkin’s lymphomas (NHL) are the fifth most common cancer type and the most common haematological malignancy. They comprise a range of genetically, phenotypically and clinically heterogeneous diseases. Most of them derived from the malignant transformation of germinal centre (GC) B cells, including DLBCL, the most common form of lymphoma. *CREBBP* loss-of-function mutations have been identified as one of the main genetic drivers involved in DLBCL [11,34]. Although ∼60% of DLBCL patients are cured with combined immnuno-chemotherapy [48], following relapse their outlook is poor with limited therapeutic options. This disease therefore remains an unmet clinical need. To expand our therapeutic arsenal and improve patient management, we must first obtain greater understanding of the molecular mechanisms underlying lymphomagenesis. The development of relevant DLBCL models which faithfully recapitulate human disease are also urgently required to test novel therapeutics.

Here, we refined our existing conditional *Crebbp* knock-out mouse models through combination with a forward insertional transposition system. This allowed for both spatial and temporal controls of *Crebbp* loss and the B-cell specific activation of the transposon GrOnc. This powerful tool led us to assess the relative contribution of the cell of origin first experiencing *Crebbp* loss and to investigate potential cooperative mutations. *Crebbp* loss within the HSPC population resulted in a very aggressive DLBCL-like disease. Mice presented with evidence of hepatosplenomegaly, as well as lymphadenopathy. Immunophenotypic characterisation revealed an aberrant B220^low^/CD19^+^ population, which infiltrated across multiple tissues. They expressed PNA and GL7 mainly, although Fas expression could also be detected in some instances. Molecularly, they activated a transcriptional program with enhanced cell cycle and cell proliferation, in part orchestrated by the up-regulation of the transcription factor E2F1 and the cell cycle regulator, cyclin D3. These genes are known to be preferentially expressed in GC B cells and contribute to the physiological clonal expansion seen during the GC reaction [49]. In contrast, transcriptomic programs related to immune cell activation, notably the CD40-CD40L signalling pathway, were silenced, preventing GC exit and terminal differentiation. Collectively, these data reflect a global rewiring of key pathways required during normal B cell maturation, most likely responsible for the malignant transformation and the disease in mice.

Although *Crebbp* loss from more committed B-cell progenitors also led to DLBCL, the latency was significantly longer. This observation was consistent with another mouse model, expressing EZH2^Y641F^ mutation following Cre expression from the Cd19 locus, which develop DLBCL after a year [50]. This suggests that the outcome of the disease varies according to the cell of origin which first acquires *Crebbp* loss-of-function mutations. When acquired in the haematopoietic stem and progenitor population, the intrinsic properties of this cell compartment (i.e, self-renewal and proliferation) would provide a favourable cellular state for transformation and potential effects on later B-cell differentiation. A later loss would also provide similar proliferative and more limited self-renewal stimuli through the GC reaction where more mature B cells will undergo active proliferation and somatic hypermutations. Alternatively, *CREBBP* mutations acquired early during B cell ontogeny would open a longer temporal window for chromatin remodelling and cell re-programming to occur. Those signals would be lacking when *CREBBP* mutations are acquired later which would explain the longer latency and the seemingly less aggressive phenotype of the disease (consistent with FL) in our model. A potential caveat of our system, given the genetic design our IM-Cd19*Crebbp*^-/-^ model, where the Cre-recombinase is knocked into the *Cd19* locus, is that CD19 haploinsufficiency may alter B-cell ontogeny, although we did not previously observe significant dysfunction [3].

Immune evasion is a well-recognised hallmark in cancer that allows tumour cells to escape immune surveillance and thus avoid macrophage-mediated phagocytosis. This phenomenon is achieved via multiple ways, such as the loss of class I MHC expression at the surface of malignant cells. Growing evidence shows that the lack of class II MHC also contributes to immune evasion in DLBCL [51,35,52]. Our data reinforces this view with a significant downregulation in the lymphoma cells of the master regulator of class II MHC gene expression *Ciita* and the subsequent downregulation of *Cd74, H2-Ab1, H2-Ob* and *H2-Eb2*, which are involved in antigen presentation and antigen processing [53]. Interestingly the B220low/CD19+ neoplastic cells also enhanced CD24 expression on their cell surface. CD24 overexpression has been reported in several solid tumours [54,55,56] and haematological malignancies [57]. In recent years, it has emerged as a novel ‘don’t eat me’ signal, which modulates anti-tumour immunity through its binding to the inhibitory receptor Siglec-10 expressed by macrophages [56]. In line with this finding, two independent studies described CD24high DLBCL as being ‘immune-cold’ with reduced expression of HLA genes and a lower abundance of tumour-infiltrating immune cells, as determined by CIBERSORT [58,59]. These data also suggest that our model may be of use in determining mechanisms of immune escape and how these might be therapeutically targeted to improve outcomes in poor risk groups such as the CD24high DLBCL patients [59].

The advent of NGS technologies has massively expanded our catalogue of somatic mutations linked to human cancer. However, our understanding of the functional implications of such mutations is still sparse and pinpointing true driver mutations remains challenging. Transposon-based screens implemented in mouse models have opened attractive avenues to overcome these limitations. We, and others, have successfully used this approach in several haematological malignancies and have identified known and novel genetic candidates [36,15]. Similarly, by transferring this technology into the B cell context, we have uncovered several common insertions sites (CIS) that have previously been associated with DLBCL and other malignancies. Globally, these CIS were enriched for specific functional categories, mainly ‘B cell development’, ‘Signalling’, ‘Cell cycle’ and ‘Transcription/Translation’. Interestingly, a similar functional observation was made from a genome-wide CRISPR screen performed on human DLBCL cell lines [42]. This highlights that the corruption of several key processes is a pre-requisite for the initiation and maintenance of transformation. Comparisons with human data also revealed significant functional overlaps, validating our system. These promising results advocate for the investigation of novel CIS that demonstrated functional relevance in our screen.

In summary, by combining *Crebbp* loss with an insertional mutagenesis system, we have developed a novel DLBCL mouse model that faithfully recapitulates well-characterised histological and molecular features of the human disease, as well as the newly described enhanced CD24 expression. From this agnostic genetic approach to identify mutational synergy in lymphoma, we uncovered an intricate regulatory network between mouse candidates and patient mutated genes. These functionally reciprocal components provide the rationale for prioritising candidates in mechanistic and therapeutic studies to improve outcomes in DLBCL.

## Acknowledgements

This study was carried out in the laboratory of B.J.P.H. with funding from Cancer Research UK (C18680/A25508, C355/A26819 and DRCRPG-Nov22/100014), MRC (MR-R009708-1), the Innovative Medicines Initiative (116026 and 945406) and the Cancer Research UK Cambridge Major Centre (C49940/A25117). This research was supported by the NIHR Cambridge Biomedical Research Centre (BRC-1215-20014), and was funded in part by the Wellcome Trust, who supported the Wellcome – MRC Cambridge Stem Cell Institute (203151/Z/16/Z) and Cambridge Institute for Medical Research (100140/Z/12/Z). The views expressed are those of the authors and not necessarily those of the NIHR or the Department of Health and Social Care. S.E.R. is supported by a Clinician Scientist Fellowship from Cancer Research UK (C67279/A27957) and a Leukaemia UK John Goldman Fellowship (2022/JGF/004). This research was supported by the CIMR Flow cytometry Core Facility. In particular, we wish to thank Drs R. Schulte and G. Grondys-Kotarba for their advice and support in cell sorting.

## Supplemental Figure Legend

**Supplementary Figure 1: Early Crebbp loss cooperates with insertional mutagenesis to significantly accelerate the B-cell lymphoma phenotype in a novel murine model**

(A). Schematic representation of the genetic systems used in the study to model conditional *Crebbp* loss upon Cre recombination and forward insertional mutagenesis based on genomic insertions of the GrOnc transposon following the expression of the Piggy Bac transposase. (B). Breeding strategy used in the study to generate the experimental mice. Only genotypes of interest are represented. (C). Schematic representation of the experimental design used in the study. 6 to 12-weeks old mice were treated with poly(I) poly(C) (pIpC) to trigger the expression of the Cre recombinase. After 6 weeks of recovery, they were challenged with SRBC injections induce PB transposase expression. Blood samples were taken regularly. Mice were culled when they showed evidence of illness. (D). Agarose gel showing *Crebbp* excision (lower band) in IM-Mx1*Crebbp*^-/-^ tissues after pIpC treatment. In contrast IM-Mx1*Crebbp*^-/-^ tissues harbour a full-length PCR product (higher band). (E). Violin plot showing the liver weight of lymphoma mice compared to control wild-type mice. Each dot represents a mouse. (F). Violin plot showing the white blood cells count of lymphoma mice compared to control wild-type mice. Each dot represents a mouse. (G-H). Violin plots showing the spleen (G) and the liver (H) weight of recipient mice after transplantation of primary tumours of indicated genotypes. Each dot represents a recipient mouse. Tum, tumour.

**Supplementary Figure 2: An aberrant B220**^**low**^**/CD19**^**+**^ **population drives a DLBCL-like disease**

(A). Violin plot showing the percentage of positive cells for the germinal centre markers PNA, GL7 and Fas assessed by flow cytometry. Cells were gated on B220^high^/CD19^+^ for the control and B220^low^/CD19^+^ for both lymphoma models. Each dot represents a mouse. (B). Histogram showing the Mean Fluorescence Intensity (MFI) of CD24 on B220^high^/CD19^+^ control cells and B220^low^/CD19^+^ lymphoma cells of indicated genotypes. (C). FACS plot showing the expression of the germinal centre markers PNA, GL7 and Fas in B220^low^/CD19^+^ cells from recipient mice after transplantation of primary tumour cells from total spleen.

**Supplementary Figure 3: Loss of CREBBP in B-cell progenitors attenuates DLBCL-like disease**

(A). Violin plot showing the liver weight of lymphoma mice of indicated genotypes compared to control wild-type mice. Each dot represents a mouse. (B). Violin plot showing the white blood cells count of lymphoma mice of indicated genotypes compared to control wild-type mice. Each dot represents a mouse. (C). Hematoxylin and eosin staining on mouse tissue sections from IM-Cd19*Crebpp*^-/-^ lymphoma models. (D-F) Violin plots showing the spleen (D) and liver (E) weight and the white blood cell count (F) of recipient mice after transplantation of IM-Cd19*Crebpp*^-/-^ primary tumours cells from total spleen. Tum, tumour.

**Supplementary Figure 4: Identification of transposon common integration sites (CIS)**

(A). Functional annotations showing the KEGG pathways for the genes insertionally targeted in mouse lymphoma models.

**Supplementary Figure 5: Positive correlation of murine models with human data**

(A). Circos plot showing the overlap between ontology terms related to the most commonly mutated genes in DLBCL patients and the commonly targeted genes in our DLBCL mouse models. (B). Networks showing the genes involved the BCR signalling pathway and the process of leukocyte differentiation. Genes are color-coded by species.

