## Supplementary material for "Mutational synergy with *CREBBP* loss in lymphomagenesis identified through forward insertional mutagenesis in a new DLBCL mouse model": Supplentary_Figures

Supplementary Table 1: List of Flow cytometry antibodies.

| Target | Fluorophore | Supplier | Clone |
|---|---|---|---|
| m/h B220 (CD45R) | APC | BioLegend | RA3-6B2 |
| m/h B220 (CD45R) | APC/Cy7 | BioLegend | RA3-6B2 |
| m B220 (CD45R) | PE/Cy7 | Biolegend | RA3-6B2 |
| m CD4 | PE | BioLegend | GK1.5 |
| m CD5 | APC | BioLegend | 53-7.3 |
| m/h CD11b | FITC | BioLegend | M1/70 |
| m CD19 | APC/Cy7 | BioLegend | 6D5 |
| m CD19 | PE | BioLegend | 6D5 |
| m CD19 | PE/Cy7 | BioLegend | 6D5 |
| m CD95 (Fas) | PE | BioLegend | SA367H8 |
| m/h GL7 | Alexa Fluor 647 | BioLegend | GL7 |
| m Gr1 (Ly6/Ly6C) | Pacific Blue | BioLegend | RB6-8C5 |
| m Ter119 | APC/Cy7 | BioLegend | TER-119 |
| PNA | Fluorescein | Vetor Laboratories | NA |

Supplementary Table 2: Consensus sequences of the top 3 clones per mouse.

| Group | Mouse ID | Clone size (n seqs) | Clone rank | V-DOMAIN Functionality | V-GENE and allele | V-REGION identity % | V-REGION identity nt | J-GENE and allele | J-REGION identity % | J-REGION identity nt |
|---|---|---|---|---|---|---|---|---|---|---|
| IM-Mx1*Crebbp*$^{-/-}$ | 7852 | 267 | 1 | productive | Musmus IGHV1-61*01 F | 96.99 | 258/266 nt | Musmus IGHJ2*01 F | 83.33 | 40/48 nt |
| | | 262 | 2 | productive | Musmus IGHV1-55*01 F | 100 | 260/260 nt | Musmus IGHJ1*03 F | 96 | 48/50 nt |
| | | 226 | 3 | productive | Musmus IGHV3-8*01 F | 100 | 260/260 nt | Musmus IGHJ4*01 F | 100 | 54/54 nt |
| | 7871 | 4979 | 1 | productive | Musmus IGHV1-55*01 F | 100 | 265/265 nt | Musmus IGHJ4*01 F | 90.74 | 49/54 nt |
| | | 1547 | 2 | productive | Musmus IGHV1-61*01 F | 100 | 266/266 nt | Musmus IGHJ4*01 F | 89.8 | 44/49 nt |
| | | 913 | 3 | productive | Musmus IGHV1-72*01 F | 100 | 266/266 nt | Musmus IGHJ2*01 F | 91.67 | 44/48 nt |
| | 8141 | 4320 | 1 | unproductive | Musmus IGHV3-8*01 F | 100 | 262/262 nt | Musmus IGHJ4*01 F | 100 | 54/54 nt |
| | | 1230 | 2 | productive | Musmus IGHV1-55*01 F | 100 | 265/265 nt | Musmus IGHJ1*03 F | 96.23 | 51/53 nt |
| | | 487 | 3 | productive | Musmus IGHV13-2*01 F | 98.52 | 267/271 nt | Musmus IGHJ3*01 F | 93.75 | 45/48 nt |
| | 8403 | 1456 | 1 | productive | Musmus IGHV1-50*01 F | 99.62 | 264/265 nt | Musmus IGHJ2*01 F | 95.83 | 46/48 nt |
| | | 1149 | 2 | unproductive | Musmus IGHV7-3*01 F | 100 | 270/270 nt | Musmus IGHJ3*01 F | 97.92 | 47/48 nt |
| | | 1047 | 3 | productive | Musmus IGHV3-1*01 F | 100 | 264/264 nt | Musmus IGHJ4*01 F | 94.44 | 51/54 nt |
| | 8635 | 4116 | 1 | productive | Musmus IGHV1-78*01 F | 100 | 265/265 nt | Musmus IGHJ1*03 F | 96.23 | 51/53 nt |
| | | 756 | 2 | productive | Musmus IGHV1-64*01 F | 99.24 | 260/262 nt | Musmus IGHJ2*01 F | 91.67 | 44/48 nt |
| | | 556 | 3 | productive | Musmus IGHV12-3*01 F | 100 | 265/265 nt | Musmus IGHJ1*03 F | 88.68 | 47/53 nt |
| IM-Mx1*Crebbp*$^{-/-}$ | 7954 | 1412 | 1 | unproductive | Musmus IGHV5-2*01 F | 100 | 265/265 nt | Musmus IGHJ1*03 F | 86.79 | 46/53 nt |
| | | 1372 | 2 | productive | Musmus IGHV3-8*01 F | 98.1 | 258/263 nt | Musmus IGHJ2*01 F | 93.75 | 45/48 nt |
| | | 1341 | 3 | productive | Musmus IGHV11-2*01 F | 100 | 265/265 nt | Musmus IGHJ1*03 F | 100 | 53/53 nt |
| | 7997 | 1581 | 1 | productive | Musmus IGHV3-8*01 F | 98.1 | 258/263 nt | Musmus IGHJ2*01 F | 93.75 | 45/48 nt |
| | | 1074 | 2 | productive | Musmus IGHV3-1*01 F | 100 | 265/265 nt | Musmus IGHJ1*03 F | 96.23 | 51/53 nt |
| | | 1009 | 3 | productive | Musmus IGHV3-8*01 F | 100 | 263/263 nt | Musmus IGHJ2*01 F | 100 | 48/48 nt |
| | 8084 | 3854 | 1 | unproductive | Musmus IGHV3-8*01 F | 100 | 262/262 nt | Musmus IGHJ4*01 F | 98.15 | 53/54 nt |
| | | 1486 | 2 | productive | Musmus IGHV3-8*01 F | 98.1 | 258/263 nt | Musmus IGHJ2*01 F | 93.75 | 45/48 nt |
| | | 1094 | 3 | productive | Musmus IGHV3-1*01 F | 100 | 265/265 nt | Musmus IGHJ1*03 F | 96.23 | 51/53 nt |
| | 8142 | 1295 | 1 | productive | Musmus IGHV3-8*01 F | 98.1 | 258/263 nt | Musmus IGHJ2*01 F | 93.75 | 45/48 nt |
| | | 910 | 2 | productive | Musmus IGHV3-1*01 F | 100 | 265/265 nt | Musmus IGHJ1*03 F | 96.23 | 51/53 nt |
| | | 884 | 3 | productive | Musmus IGHV3-8*01 F | 100 | 263/263 nt | Musmus IGHJ2*01 F | 100 | 48/48 nt |
| | 8337 | 1693 | 1 | productive | Musmus IGHV3-8*01 F | 98.1 | 258/263 nt | Musmus IGHJ2*01 F | 93.75 | 45/48 nt |
| | | 1206 | 2 | productive | Musmus IGHV3-1*01 F | 100 | 266/266 nt | Musmus IGHJ1*03 F | 96.23 | 51/53 nt |
| | | 979 | 3 | productive | Musmus IGHV3-8*01 F | 98.48 | 259/263 nt | Musmus IGHJ2*01 F | 93.75 | 45/48 nt |
| | 8354 | 1035 | 1 | productive | Musmus IGHV3-8*01 F | 98.1 | 258/263 nt | Musmus IGHJ2*01 F | 93.75 | 45/48 nt |
| | | 906 | 2 | productive | Musmus IGHV3-8*01 F | 100 | 262/262 nt | Musmus IGHJ2*01 F | 100 | 48/48 nt |
| | | 717 | 3 | productive | Musmus IGHV3-1*01 F | 100 | 264/264 nt | Musmus IGHJ1*03 F | 96.23 | 51/53 nt |
| | 8583 | 924 | 1 | productive | Musmus IGHV3-8*01 F | 98.09 | 257/262 nt | Musmus IGHJ2*01 F | 93.75 | 45/48 nt |
| | | 665 | 2 | productive | Musmus IGHV3-1*01 F | 100 | 266/266 nt | Musmus IGHJ1*03 F | 96.23 | 51/53 nt |
| | | 583 | 3 | productive | Musmus IGHV3-8*01 F | 100 | 263/263 nt | Musmus IGHJ2*01 F | 100 | 48/48 nt |

Supplementary Table 3: List of genes with at least 3 independent insertions.

| Mouse Ensembl ID | Mouse gene name | No of insertions |
|---|---|---|
| ENSMUSG00000057098 | Ebf1 | 29 |
| ENSMUSG00000020919 | Stat5b | 24 |
| ENSMUSG00000042817 | Flt3 | 24 |
| ENSMUSG00000027852 | Nras | 20 |
| ENSMUSG00000024789 | Jak2 | 18 |
| ENSMUSG00000014030 | Pax5 | 16 |
| ENSMUSG00000044303 | Cdkn2a | 16 |
| ENSMUSG00000024241 | Sos1 | 14 |
| ENSMUSG00000024785 | Rcl1 | 10 |
| ENSMUSG00000092329 | Gm20388 | 8 |
| ENSMUSG00000005268 | Prlr | 7 |
| ENSMUSG00000024817 | Uhrf2 | 7 |
| ENSMUSG00000002107 | Celf2 | 6 |
| ENSMUSG00000024782 | Ak3 | 6 |
| ENSMUSG00000035649 | Zcchc7 | 6 |
| ENSMUSG00000038658 | Ric1 | 6 |
| ENSMUSG00000055932 | Fto | 6 |
| ENSMUSG00000004698 | Hdac9 | 5 |
| ENSMUSG00000005886 | Ncoa2 | 5 |
| ENSMUSG00000018654 | Ikzf1 | 5 |
| ENSMUSG00000026872 | Zeb2 | 5 |
| ENSMUSG00000030067 | Foxp1 | 5 |
| ENSMUSG00000036550 | Cnot1 | 5 |
| ENSMUSG00000040929 | Rfx3 | 5 |
| ENSMUSG00000052920 | Prkg1 | 5 |
| ENSMUSG00000052942 | Glis3 | 5 |
| ENSMUSG00000054611 | Kdm2a | 5 |
| ENSMUSG00000064202 | 4430402I18Rik | 5 |
| ENSMUSG00000085936 | 2610307P16Rik | 5 |
| ENSMUSG00000097855 | A930007I19Rik | 5 |
| ENSMUSG00000013663 | Pten | 4 |
| ENSMUSG00000019982 | Myb | 4 |
| ENSMUSG00000021892 | Sh3bp5 | 4 |
| ENSMUSG00000022346 | Myc | 4 |
| ENSMUSG00000022637 | Cblb | 4 |
| ENSMUSG00000024420 | Zfp521 | 4 |
| ENSMUSG00000025626 | Phf6 | 4 |
| ENSMUSG00000026395 | Ptprc | 4 |
| ENSMUSG00000026721 | Rabgap1l | 4 |
| ENSMUSG00000026737 | Pip4k2a | 4 |
| ENSMUSG00000028163 | Nfkb1 | 4 |
| ENSMUSG00000031453 | Rasa3 | 4 |
| ENSMUSG00000032035 | Ets1 | 4 |
| ENSMUSG00000037138 | Aff3 | 4 |
| ENSMUSG00000040451 | Sgms1 | 4 |
| ENSMUSG00000045333 | Zfp423 | 4 |
| ENSMUSG00000046138 | 9930021J03Rik | 4 |
| ENSMUSG00000053477 | Tcf4 | 4 |
| ENSMUSG00000115520 | AC119177.2 | 4 |
| ENSMUSG00000000861 | Bcl11a | 3 |
| ENSMUSG00000004980 | Hnrnpa2b1 | 3 |
| ENSMUSG00000021365 | Nedd9 | 3 |
| ENSMUSG00000022353 | Mtss1 | 3 |
| ENSMUSG00000022708 | Zbtb20 | 3 |
| ENSMUSG00000022885 | St6gal1 | 3 |
| ENSMUSG00000024251 | Thada | 3 |
| ENSMUSG00000024780 | Cdc37l1 | 3 |
| ENSMUSG00000026288 | Inpp5d | 3 |
| ENSMUSG00000028121 | Bcar3 | 3 |
| ENSMUSG00000028525 | Pde4b | 3 |
| ENSMUSG00000028920 | Fbxo42 | 3 |
| ENSMUSG00000030452 | Nipa2 | 3 |
| ENSMUSG00000031668 | Eif2ak3 | 3 |
| ENSMUSG00000032253 | Phip | 3 |
| ENSMUSG00000032393 | Dpp8 | 3 |
| ENSMUSG00000033991 | Ttc37 | 3 |
| ENSMUSG00000035473 | Galm | 3 |
| ENSMUSG00000036748 | Cuedc2 | 3 |
| ENSMUSG00000038151 | Prdm1 | 3 |
| ENSMUSG00000038481 | Cdk19 | 3 |
| ENSMUSG00000039943 | Plcb4 | 3 |
| ENSMUSG00000040274 | Cdk6 | 3 |
| ENSMUSG00000041112 | Elmo1 | 3 |
| ENSMUSG00000044471 | Lncpint | 3 |
| ENSMUSG00000048612 | Myof | 3 |
| ENSMUSG00000053007 | Creb5 | 3 |
| ENSMUSG00000055447 | Cd47 | 3 |
| ENSMUSG00000073563 | Csnk1g3 | 3 |
| ENSMUSG00000075014 | Gm10800 | 3 |
| ENSMUSG00000078970 | Wdr92 | 3 |
| ENSMUSG00000079509 | Zfx | 3 |
| ENSMUSG00000089862 | Umad1 | 3 |
| ENSMUSG00000097640 | Gm20033 | 3 |
| ENSMUSG00000102805 | Gm37240 | 3 |
| ENSMUSG00000108563 | Gm44686 | 3 |
| ENSMUSG00000111200 | Gm33728 | 3 |
| ENSMUSG00000115852 | AC169509.1 | 3 |

Supplementary Figure 1

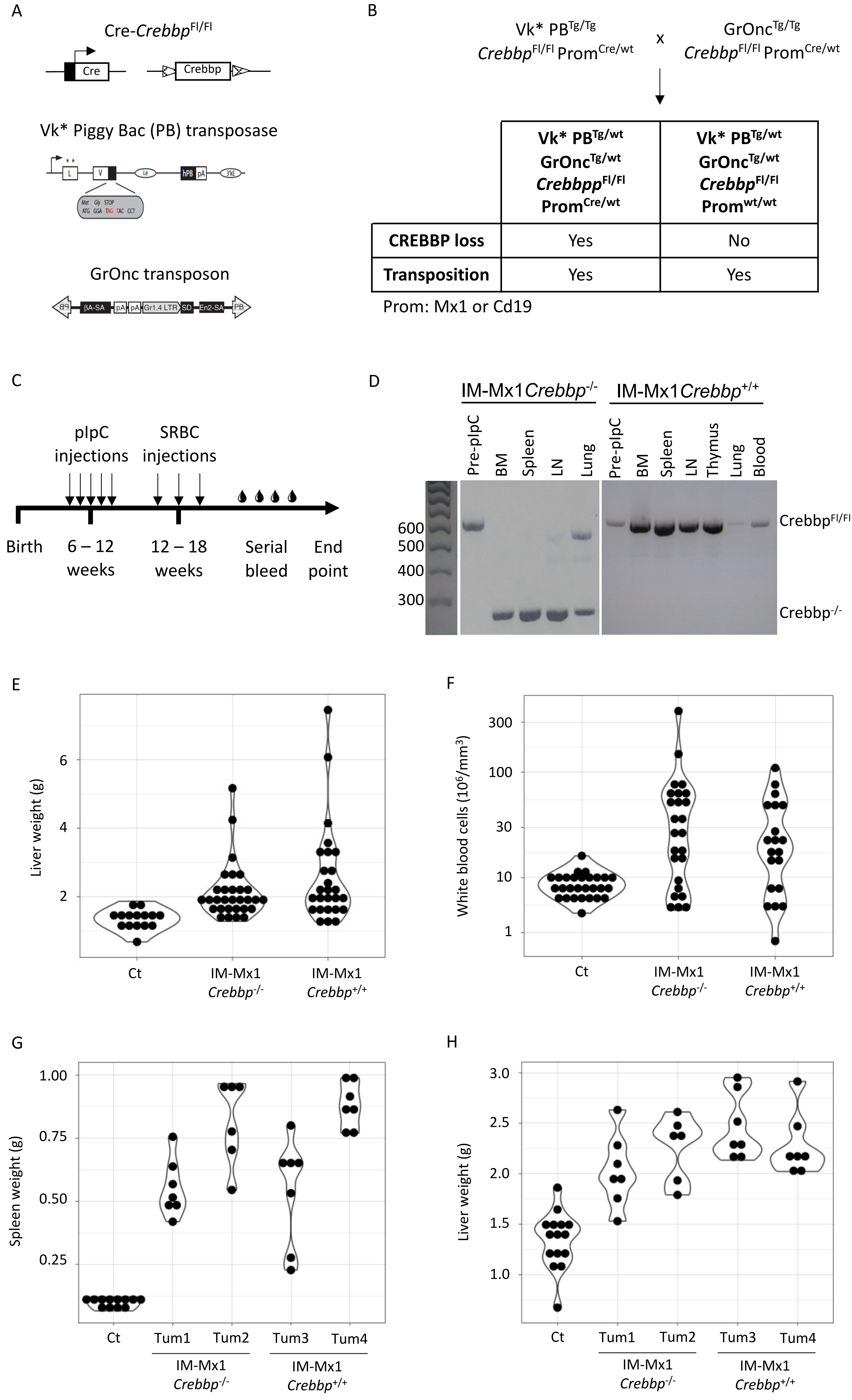

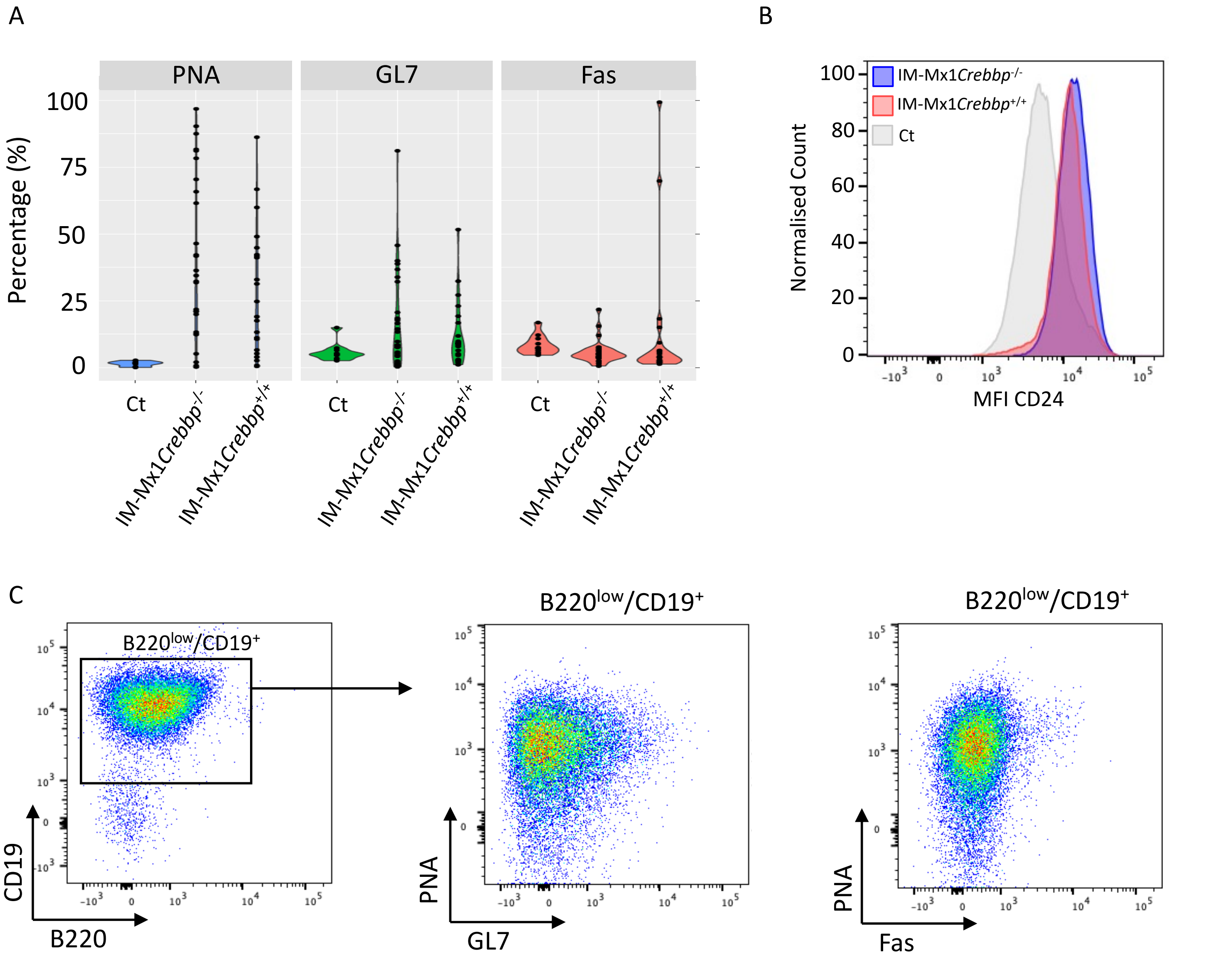
A
PNA
GL7
Fas
Percentage (%)
100
75
50
25
0
Ct
IM-Mx1Crebbp-/-
IM-Mx1Crebbp+/+
B
IM-Mx1Crebbp-/-
IM-Mx1Crebbp+/+
Ct
Normalised Count
MFI CD24
C
B220low/CD19+
CD19
B220
PNA
GL7
Fas

Supplementary Figure 3

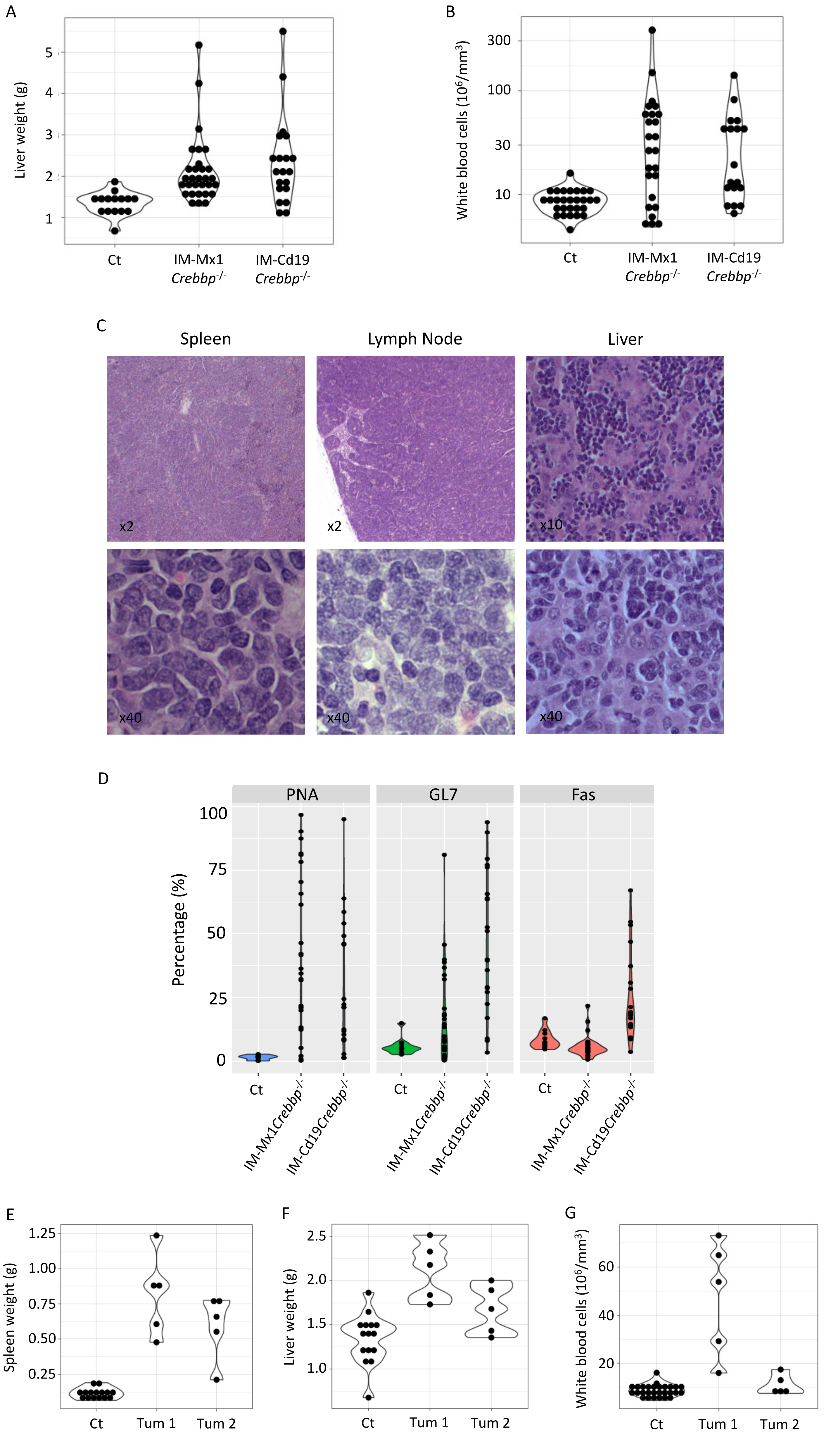

A

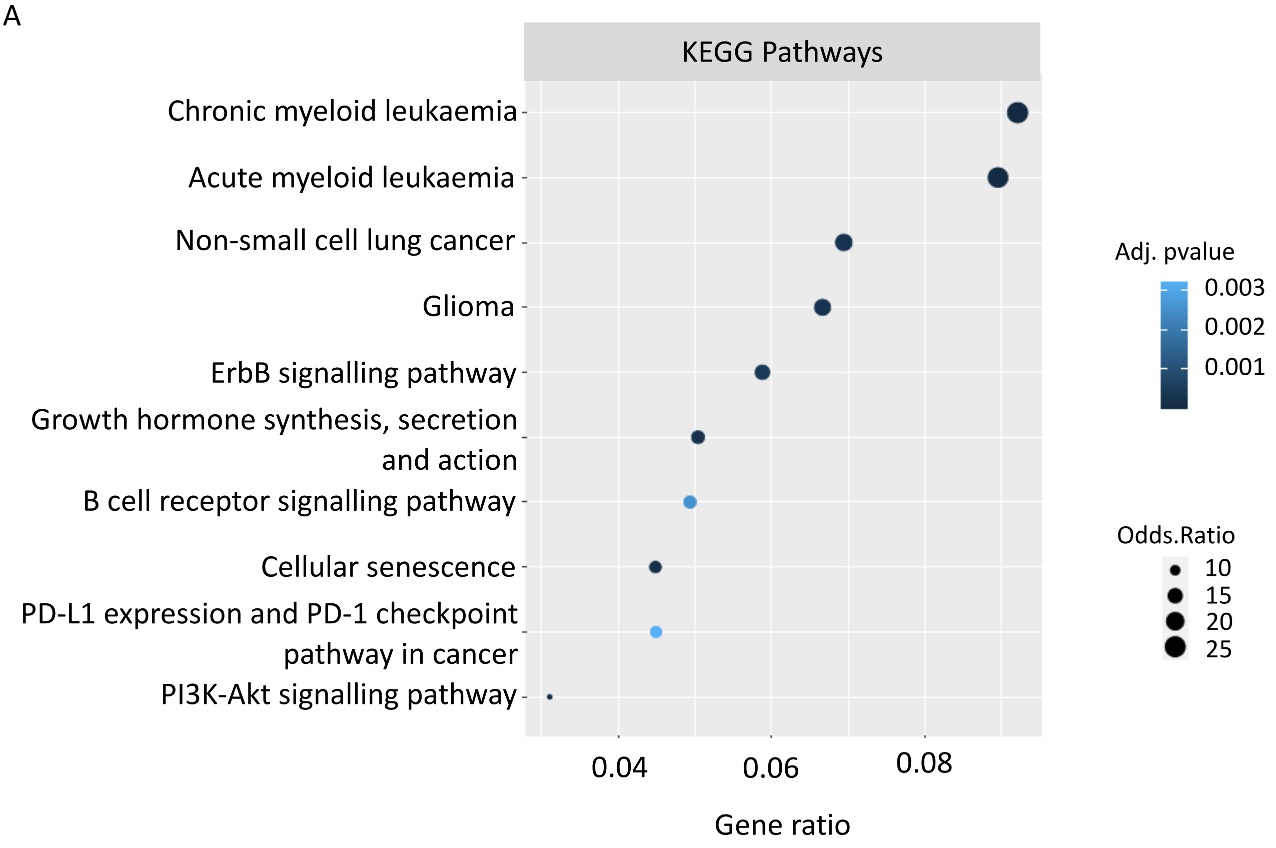
KEGG Pathways
Chronic myeloid leukaemia
Acute myeloid leukaemia
Non-small cell lung cancer
Glioma
ErbB signalling pathway
Growth hormone synthesis, secretion and action
B cell receptor signalling pathway
Cellular senescence
PD-L1 expression and PD-1 checkpoint pathway in cancer
PI3K-Akt signalling pathway
0.04
0.06
0.08
Gene ratio
Adj. pvalue
0.003
0.002
0.001
Odds.Ratio
10
15
20
25

Supplementary Figure 5

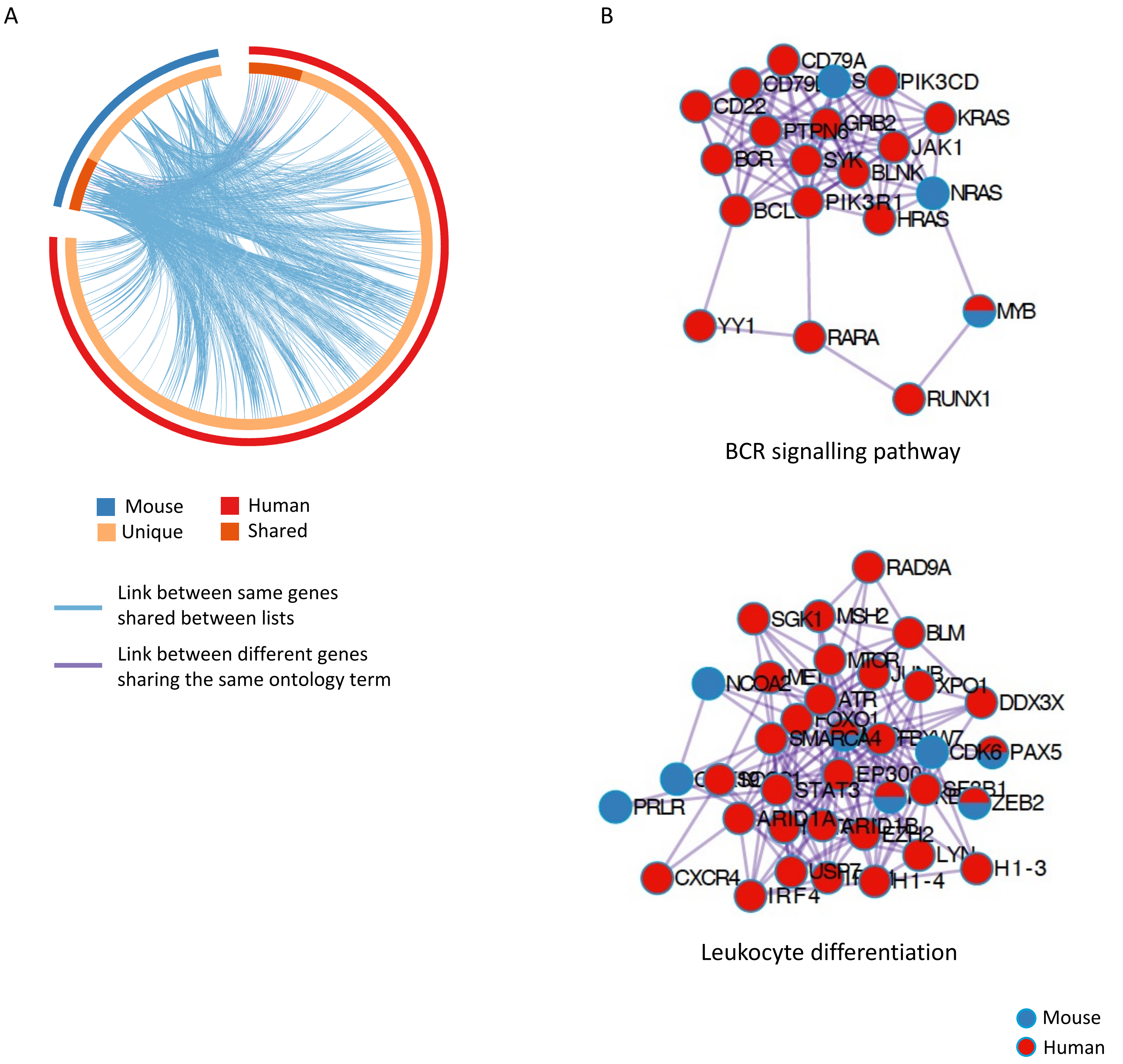
